# Oblique light-sheet tomography: fast and high resolution volumetric imaging of mouse brains

**DOI:** 10.1101/132423

**Authors:** Arun Narasimhan, Kannan Umadevi Venkataraju, Judith Mizrachi, Dinu F. Albeanu, Pavel Osten

## Abstract

Present light sheet fluorescence microscopes lack the wherewithal to image the whole brain (large tissues) with uniform illumination/detection parameters and high enough resolution to provide an understanding of the various aspects of neuroanatomy. To overcome this, we developed an oblique version of the light sheet microscope (Oblique Light Sheet Tomography, OLST) that includes a high magnification objective and serial sectioning, for volumetric imaging of the whole mouse brain at high spatial resolution at constant illumination/detection. We developed a novel gelatin based re-embedding procedure that makes the cleared brain rigid so that it can sectioned using our integrated microtome. Here, we characterize OLST and show that it can be used to observe dendritic morphology, spines and follow axons over a few mm in the mouse brain.

## INTRODUCTION

Imaging the whole-brain in high resolution helps provide information regarding the cell type distribution, dendritic morphology and long-range projection. Light microscopy provides fast and high-resolution images to understand the mesoscale connectivity and the micro-circuitry within the brain^1^. Serial two-photon tomography (STPT)^2-5^ achieved full volumetric brain imaging at constant and high spatial resolution by integrating laser scanning two-photon microscopy with vibratome-based tissue sectioning: after imaging the top of the tissue, this region is mechanically removed by the integrated vibratome and the process repeats until the entire brain is imaged. However, one limitation this method is its slow data acquisition time: for example, imaging one mouse brain at 0.45*0.45*1.33 μm takes 8 days using the most advanced STPT instrumentation^5^. Alternatively, fMOST^6^ and WVT^7^ acquire high-resolution whole brain complete data within 18 or 3 days. There exists a need to do whole brain imaging at high spatial resolution with high speed to uncover the morpholology and long-range connectivity pattern.

Light sheet fluorescence microscopy (LSFM)^8^, which combines illumination of a cleared brain sample by a thin light sheet and wide-field orthogonal imaging by a fast sCMOS camera, achieved fast and high resolution volumetric imaging for samples that are few hundred microns wide, such as the embryogenesis of *Drosophila melanogaster^9^* and larval zebrafish^10^,^11^. However scattering limits the use of light sheet imaging for large tissues, such as a mouse or a rat brain. A critical part of LSFM for large tissue volumetric imaging, such as a mouse or a rat brain, is the step of reducing light scattering by removing lipids in the brain using either organic^12^,^13^,^14^ or aqueous based solutions^15^, ^16-18^, before this delipidated tissue is immersed in an refractive index (RI) matching solution, to make it optically transparent^19^. These improvements have reduced the scattering and increased the mean free path of the photons, the size of the tissue (few mm) and the extent of clearing (non-uniform) and the depth of imaging (ballistic scattering losses and collection efficiency) limit the imaging quality.

Typical light sheet microscopes use a low magnification objective (<5x) to image the whole brain of the mouse^12^,^13^,^20^,^24^. This approach has been used to probe c-fos activation patterns^12^ or follow projections of dense axon bundles^25^, but does not have the resolving power to view dendritic morphology or follow long range axons that form an important part of neuro-anatomy and provide understanding of the functioning of the brain. High magnification objectives have been used to image only portion of the brain such as the hippocampus^20^ or the cerebellum^26^ as the working distance of the objective limits the depth of imaging within the tissue. Clarity-optimized light-sheet microscopy (COLM)^18^, another version of LSFM that uses specially developed extra-long working distance 10x (0.6 NA, 8mm WD) objectives to image the whole brain or use a 25x (0.95 NA, 8mm WD) to image a portion of the brain. While COLM can image the whole brain of the mouse, it suffers from image quality degradation at higher depths leading to spherical aberrations and decreased SNR due to inefficient sample clearing and the density gradient within tissue (e.g. white vs. gray matter)^19^. Modeling spherical aberration and deconvolution^27^ can in principle minimize these issues, but require *a priori* knowledge and/or iterative calculation of RI and scattering for each imaged location that can vary widely across brain regions and samples. Adaptive optics^28^, 0-scanning^22^ and structured illumination^29^ have also been proposed for correcting aberrations and increasing image quality, but these techniques suffer from substantial increases in acquisition time thereby reducing the advantages of LSFM as a fast volumetric imaging modality.

Here we present a novel LSFM-based imaging method, named oblique light-sheet tomography (OLST), that achieves fast imaging of whole mouse brains with high spatial resolution and near-constant imaging conditions throughout the entire sample. In OLST, the sample is imaged from the top using oblique LSFM illumination and detection strategy^30-33^ and the imaged top portion of the tissue is removed by vibratome sectioning, much in the same way as in STPT (Fig 1a). The new method thus combines the best of both worlds—optimal and near-constant imaging conditions achieved by serial sectioning and fast data acquisition achieved by LSFM with sCMOS camera. Here, we demonstrate the unique capabilities of OLST by acquiring whole mouse brain dataset at 0.75*0.75*2.5μm resolution in approximately 14 hrs. We show that these high resolution images with high SNR acquired from the OLST can be used to reconstruct the dendritic morphology of individual neurons and also can be used to do long range tracing of individual neurons.

## RESULTS

### Oblique Light Sheet Tomography (OLST)

The development of OLST includes the OLST instrument, data acquisition and analysis software, and an optimized brain clearing procedure. The instrument’s imaging core (Fig. 1a, b and c) consists of two orthogonal non-symmetric water-dipping objectives: a 10x Nikon PlanFluor (0.3 NA, 3.5mm WD, effective NA 0.065) used for sample illumination and a 16x Nikon LWD (0.8 NA, 3mm WD) used as the detection objective (Fig. 1b and c). The light sheet (488 or 561nm) was generated by digital scanning of a small laser beam using x-galvanometric scanner that was placed in the conjugate back focal plane of the illumination objective. The light sheet parameters were 5.8 μm and 20.1μm at the center and edge of the acquisition (Supplementary Fig. 1) respectively. The emitted fluorescence from the sample was collected using a 16x Nikon objective that was coupled to a 200mm tube lens and imaged onto the Andor Zyla 4.2 sCMOS camera. The pixel size is given by 0.41*0.41μm while the PSF of the microscope was measured as 0.75*0.75μm along x and y, and 6.9um along the detection axis (Supplementary Fig. 2). We typically read out only one half of the sCMOS chip corresponding to the central (flatter and thinner) portion of the Gaussian light sheet. This allows for imaging the surface of the brain (oblique depth ~400 μm) in an XY raster pattern (Fig. 1a) using the motorized stages. Once the raster scan is completed, the brain is moved to an integrated vibratome to section the imaged the top 250μm thick layer of tissue. The sequence of raster scans and automated sectioning was repeated across the whole brain (Fig. 1a). To acquire at the fastest speed (100Hz), we do not use the virtual slit/light sheet mode of the camera and we move the sample continuously at 0.5mm/s or 0.25mm/s. Using OLST, we acquired a whole brain high-resolution data at 0.75*0.75*5m or 0.75*0.75*2.5μm data in 8 hrs or 14 hrs respectively.

**Figure 1.**
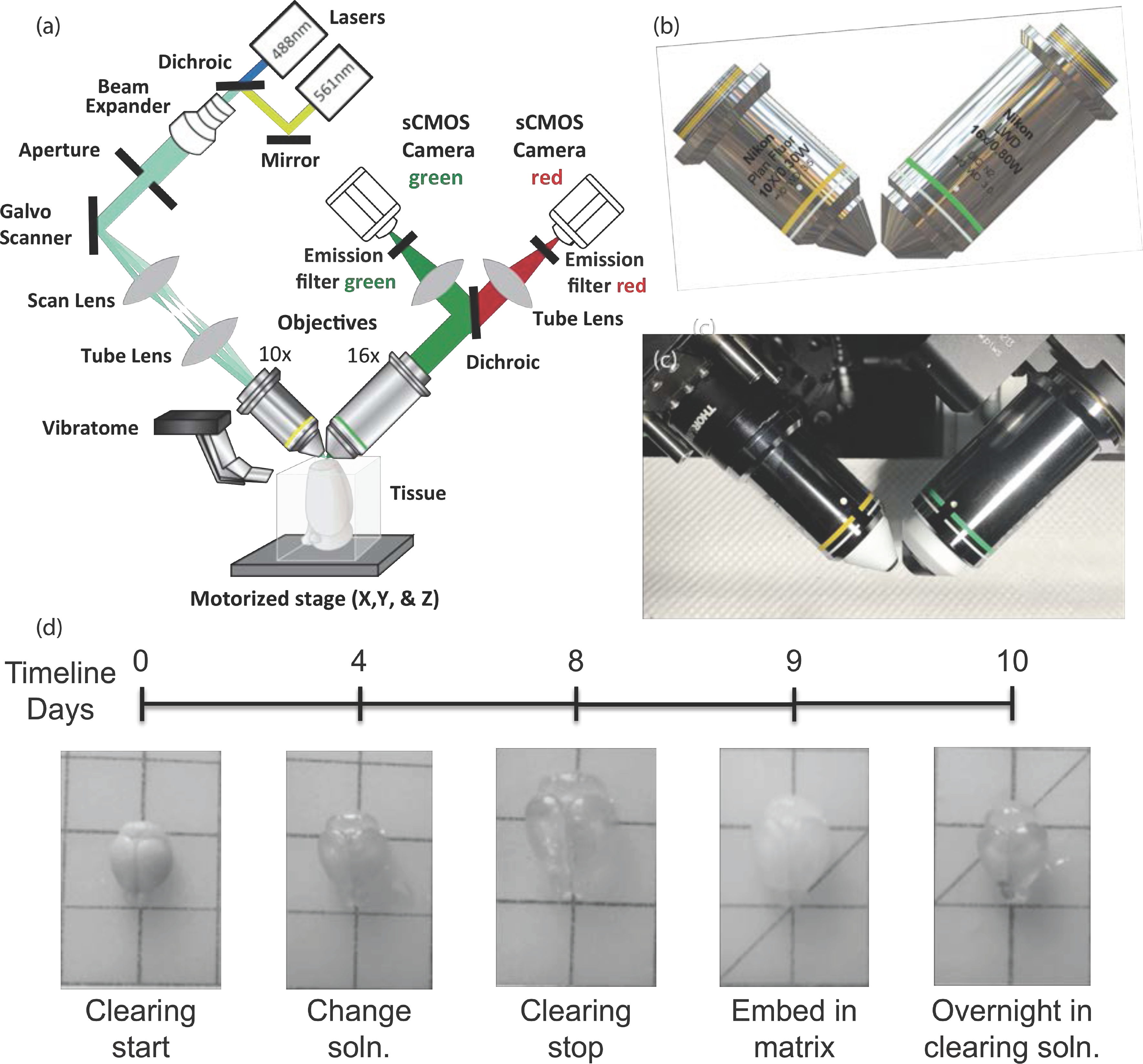
(a) Oblique Light Sheet Tomography schematic consisting of the lasers, galvanometric scanner, beam expanders, lens, illumination and detection objectives, filters, sCMOS cameras, the vibratome, the cleared mouse brain on 3-axis motorized stage. (b) CAD rendering of the two non-symmetric objectives used for illumination (10x) and the detection (16x). (c) Zoom view of the actual system that consists of the Nikon 10x illumination and Nikon 16x detection objectives, respectively. (d) mCUBIC clearing protocol and timeline-a fixed brain incubated in clearing solution at 37°C, after 4 days the solution is changed. After 1 week, the clearing is stopped and the brain is washed PB. The following day, the brain is embedded in 5% gelatin matrix to make it rigid enough to be sectioned by a microtome. Prior to imaging, the gelatin embedded brain is incubated in clearing solution (37°C) to make the brain translucent.

The sample for OLST imaging needs to be chemically cleared and, in addition, remain rigid to allow for good mechanical sectioning. To this end, we have optimized the previously published CUBIC method^16^,^17^. We cleared a mouse brain using the newly optimized mCUBIC (modified CUBIC) clearing solution that consists of 25% w/w N,N,N’,N’-tetrakis(2-hydroxypropyl)ethylenediamine, 15% w/w of Triton-X and 60% w/w of dH2O (Fig. 1d). Briefly, we immersed a perfused brain in the mCUBIC clearing solution and incubated at 37C in a shaker oven. After 4 days, we changed the clearing solution (Fig. 1d). At day 8, we stopped the clearing process by washing off the clearing solution using 0.05M PB solution. This washing step reduced the size of the brain after which we embedded the brain in 5% gelatin matrix and crosslinked overnight in 4% PFA. Prior to imaging in OLST, we incubated the cleared-gelatin embedded brain in mCUBIC clearing solution overnight. Now, we prepared an agarose block with the brain in it to help maintain the position constant while imaging and sectioning in OLST (Fig. 1b and c). This clearing and gelatin embedding procedure made the brain translucent for imaging in OLST and rigid enough to be sectioned using our integrated vibratome.

### OLST enables whole brain imaging with uniform illumination parameters

To demonstrate that OLST can achieve whole brain near-constant illumination/detection without photobleaching effects, we imaged a cleared/gelatin-embedded GAD-H2B-GFP mouse brain (Fig. 2), a mouse line that labels the entire GABAergic population of neurons with a nuclear GFP tag, in OLST with a step size of 5 μm. An example coronal region within the brain is shown in Fig 2d. Fig. 2e, f, g and h show zoomed in regions at different locations within the example coronal section of Fig. 2d. As our lateral pixel resolution (0.41*0.41μm) is high, we can clearly observe individual nuclei even in densely expressed regions of the brain. Since, the illumination/detection in OLST is oblique with respect to the tissue surface, illumination/detection conditions are uniform across the entire lateral field of view, so there is no need for iterative focusing^27^, dual sided illumination or multi-view imaging^10^. Further, photobleaching is minimal as seen in an example section (Supplementary Fig. 3) about 2.5mm posterior to the one shown in Fig 2d. The size of the tissue nor the working distance of the high magnification objectives do not limit us as the integrated vibratome sections out the imaged tissue volume. Any fluorescent tissue that is cleared and gelatin-embedded can be imaged in our OLST with micrometric resolution.

**Figure 2.**
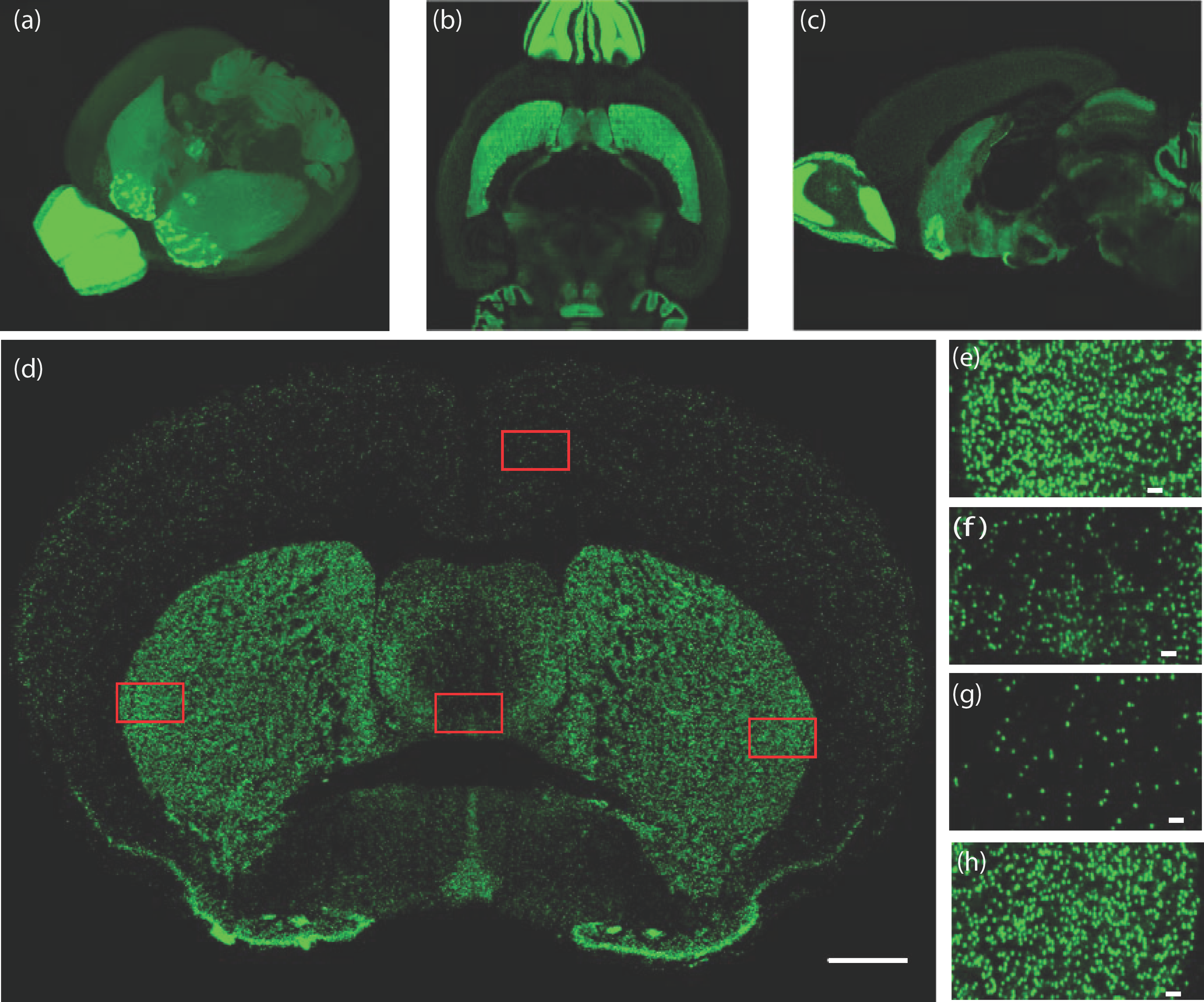
(a) Low resolution volume rendering of the GADH2B-GFP whole brain (b) Horizontal and (c) Sagittal sections from the whole brain. (d) A representative coronal section from the GADH2B-GFP. Scale bar-1mm. (e), (f), (g) and (h) zoom in view of the same section within the various regions of the brain, lateral left hemisphere (e), medial ventral portion (f), medial dorsal area (g) and lateral right hemisphere (h). Scale bar-50μm.

### OLST images the whole brain at high-spatial resolution

An important goal of neuroanatomy is to reconstruct the anatomy of individual neurons in various levels such as the dendritic morphology, axonal projections and spines. To show that OLST can be used to reconstruct dendritic morphology, observe spines and follow long range axons, a Thy1-GFP mouse brain was imaged using the OLST. We used a 2.5μm x-step to reduce the motion artifacts and to resolve smaller structures, such as spines, morphology from whole brain. The whole brain was imaged in 14hrs (~11TB raw data, including sectioning time). Fig 3 shows a representative section imaged using OLST of a Thy1-GFP(+) mouse. We can see spines in the along the whole brain (Fig 3b, c and d). OLST is the first microscope where we can image a whole-mouse brain at spine level resolution.

**Figure 3.**
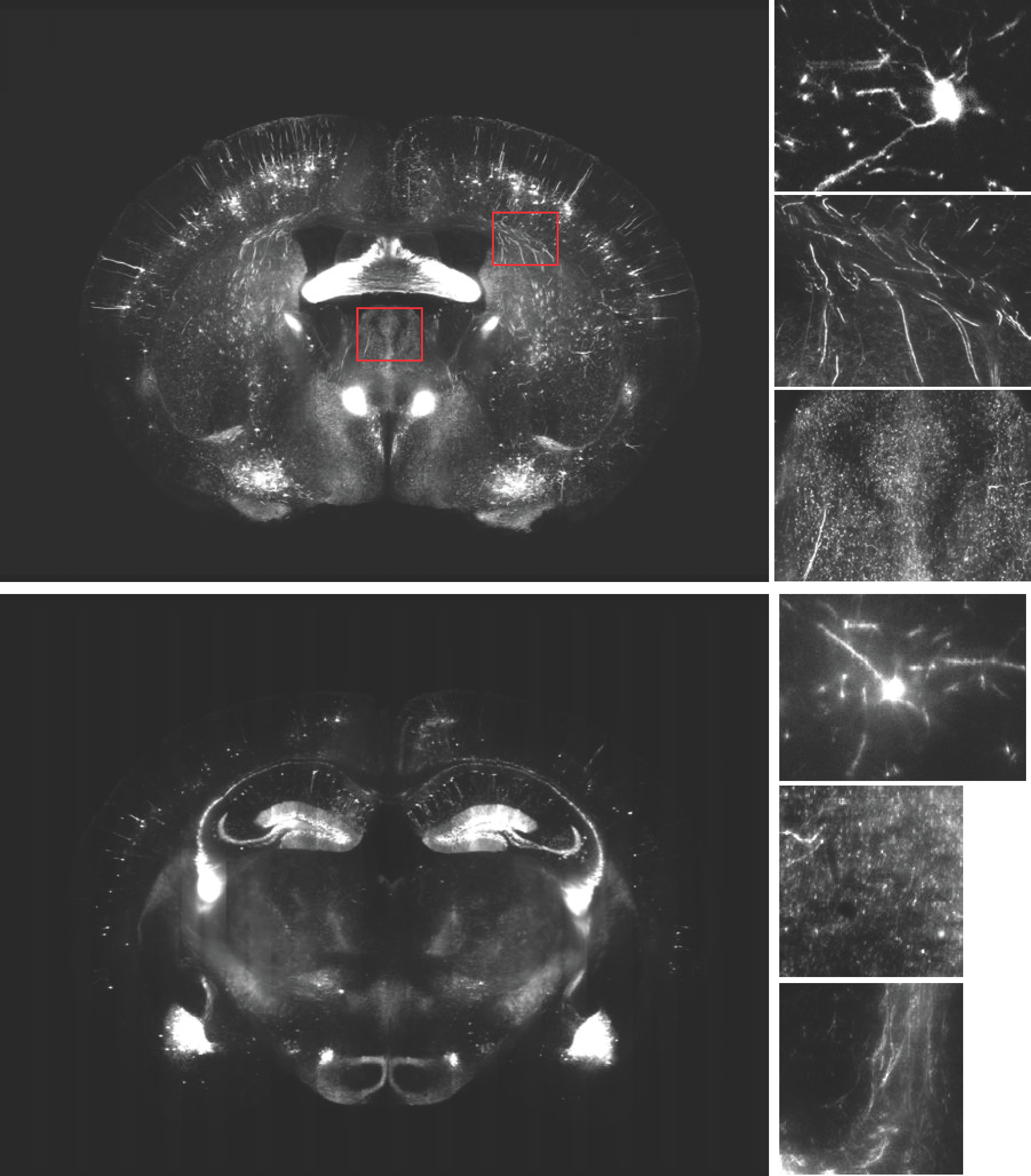
(a) and (b) Sagittal and Horizontal rendering of a Thy1-GFP brain imaged using OLST. Scale bar-2mm. (c) and (e) Coronal images at two different regions of the brain-(c) anterior and (e) posterior. Scale bar-2mm. (d) and (f) High-resolution images from the coronal images showing spines. Scale bar-25μm.

## CONCLUSIONS

We have developed a robust light sheet fluorescence microscope in combination with serial sectioning, in the oblique geometry, that can image whole tissues at high resolution. We adapted and modified the CUBIC clearing protocol to clear the sample and then to embed it in gelatin to make it rigid enough to be sectioned using the integrated microtome. The combination of serial sectioning and LSFM modality provides us the ability to image whole brain at near-uniform illumination parameters and high speed, while the use of high magnification objectives help us to provide a high spatial resolution images along the whole brain.

## ACKNOWLEDGEMENTS

This work was supported by NIH Grant U01MH105971 to P.O. We would like to thank Keerthi Krishnan and Josh Huang for providing the GADH2B_GFP mouse line.

